# Enabling Semantic Queries Across Federated Bioinformatics Databases

**DOI:** 10.1101/686600

**Authors:** Ana Claudia Sima, Tarcisio Mendes de Farias, Erich Zbinden, Maria Anisimova, Manuel Gil, Heinz Stockinger, Kurt Stockinger, Marc Robinson-Rechavi, Christophe Dessimoz

**Affiliations:** ZHAW Zurich University of Applied Sciences, Switzerland; Department of Computational Biology, University of Lausanne, Switzerland; Center for Integrative Genomics, University of Lausanne, Switzerland; SIB Swiss Institute of Bioinformatics, Lausanne, Switzerland; Department of Ecology and Evolution, University of Lausanne, Switzerland; Department of Genetics, Evolution, and Environment, University College London, UK; Department of Computer Science, University College London, UK

**Author notes:** Equal contribution (joint first authors) (Ana Claudia Sima, Tarcisio Mendes de Farias). Correspondence: *{, }.

**Keywords:** data interoperability, ontology, SPARQL

## Abstract

**Motivation:** Data integration promises to be one of the main catalysts in enabling new insights to be drawn from the wealth of biological data available publicly. However, the heterogeneity of the different data sources, both at the syntactic and the semantic level, still poses significant challenges for achieving interoperability among biological databases.

**Results:** We introduce an ontology-based federated approach for data integration. We applied this approach to three heterogeneous data stores that span different areas of biological knowledge: 1) Bgee, a gene expression relational database; 2) OMA, a Hierarchical Data Format 5 (HDF5) orthology data store, and 3) UniProtKB, a Resource Description Framework (RDF) store containing protein sequence and functional information. To enable federated queries across these sources, we first defined a new semantic model for gene expression called GenEx. We then show how the relational data in Bgee can be expressed as a virtual RDF graph, instantiating GenEx, through dedicated relational-to-RDF mappings. By applying these mappings, Bgee data are now accessible through a public SPARQL endpoint. Similarly, the materialised RDF data of OMA, expressed in terms of the Orthology ontology, is made available in a public SPARQL endpoint. We identified and formally described intersection points (i.e. virtual links) among the three data sources. These allow performing joint queries across the data stores. Finally, we lay the groundwork to enable nontechnical users to benefit from the integrated data, by providing a natural language template-based search interface.

**Project URL:** http://biosoda.expasy.org, https://github.com/biosoda/bioquery

## 1. Introduction

One key promise of the postgenomic era is to gain new biological insights by integrating different types of data (e.g. 1, 2). For instance, by comparing disease phenotypes in humans with phenotypes produced by particular mutations in model species, it is possible to infer which human genes are involved in the disease (3).

A wealth of biological data is available in public data repositories; over one hundred key resources are featured in the yearly Nucleic Acids Research annual database issue (4). However, these databases vary in the way they model their data (e.g. relational, object-oriented, or graph database models), in the syntaxes used to represent or query the data (e.g. markup or structured query languages), and in their semantics. This heterogeneity poses challenges to integrating data across different databases.

Ontologies have been widely used to achieve data integration and semantic data exchange (5–12). In this paper, by ontology, we adopt the broadly accepted definition in data and knowledge engineering of “a formal, explicit specification of a shared conceptualization” (13). The relevance of ontologies in life sciences can be illustrated by the fact that repositories such as BioPortal (14) contain more than seven hundred biomedical ontologies, and the OBO Foundry (15) more than 170 ontologies. Moreover, major life-sciences databases use ontologies to annotate and schematize data, such as UniProt (16) or ChEBI (17). Ontologies are important to enable knowledge sharing.

Currently, however, even when resources describe their data with ontologies, aligning these ontologies and combining information from different databases remain largely manual tasks, which require intimate knowledge of the way the data is organised in each source. This is despite a plethora of existing literature on data integration approaches, in particular in biological research (surveys of these approaches, as well as the challenges involved, include (18–20)). Projects such as KaBOB (21), Bio2RDF (22) and Linked Life Data (23) link different life science resources using a common ontology and data conventions. However, their centralised architecture makes it difficult to remain up-to-date and to scale up. For example, when querying the number of UniProt protein entries over the outdated and centralised Linked Life Data approach, we can only count around 10% of the 230 million entries that are in the current UniProt release (see Supplementary Material Section S1 for further explanations). To avoid this issue, federated approaches have recently been proposed (24–27), but to the best of our knowledge, none of them proposes a vocabulary and patterns to extensively, explicitly and formally describe how the data sources can be interlinked further than only considering “same as”-like mappings; in effect, they put the burden on the users to find out precisely *how* to write a conjunctive federated query. An emerging research direction entails automatically discovering links between datasets using Word Embeddings (28). We did not pursue this approach, given that it is computationally expensive and that for our study writing the relational-to-RDF mappings proved more straightforward. However, Word Embeddings would be important for the case of integrating more data sources for which the connecting links (join points) are not clearly known. Most of the existing federated approaches address the problem of multiple database models by explicitly converting and storing the data into the same type of storage engine in order to achieve data interoperability. This often implies data duplication, which complicates maintenance. Among the aforementioned federated approaches, we can highlight the approach in (26), which requires less human interventions to generate federated queries. Nonetheless, this approach was mostly designed for chemical substance data based on predefined “same as” mappings and handcrafted query patterns. Moreover, when considering the generated SPARQL query examples, they are mostly disjunctive queries (i.e. union) rather than complex conjunctive queries (i.e. intersection) — which are our main focus. As opposed to other federated systems, such as BioFed (24), we do not focus on benchmarking or improving the performance of the underlying federation engine. However, our experiments with federated queries on the integrated data corroborate existing studies in showing that federation engines exhibit significant performance degradation when processing queries that involve large intermediate result sets (29).

To address the problem of semantic, syntactic and data model heterogeneity, we propose an ontology-driven linked data integration architecture. We apply this architecture to build a system that federates three bioinformatics databases containing: evolutionary relationships among genes across species (OMA), curated gene expression data (Bgee), and biological knowledge on proteins (UniProt). In Supplementary Material, we summarise the key data provided by Bgee, OMA and UniProt (Table S3). Each of the three databases uses a different technical approach to store information: a Hierarchical Data Format 5 (HDF5, http://www.hdfgroup.org/HDF5/) data store for OMA (30); a relational database for Bgee (31); and a Resource Description Framework (RDF) store for UniProt (16). Our main contribution is to enable researchers to jointly query (i.e. conjunctive queries) the three heterogeneous databases using a common query language, by introducing and leveraging “virtual links” between the three sources. Furthermore, we show how relational data can be made interoperable with RDF data *without* requiring the original relational data to be duplicated into an RDF storage engine. This can be achieved by constructing dedicated relational-to-RDF mappings, allowing the unmodified original data to be queried via the structured query language SPARQL (32). In our proposed architecture, we illustrate this through the example of the Bgee relational database.

Moreover, for the purpose of building the federated data access system, we make the following additional contributions: (i) a semantic model for gene expression; (ii) an extension and adaptation of the Vocabulary of Interlinked Datasets (VoID) (33); (iii) public SPARQL 1.1 (32) query endpoints for OMA and Bgee; and (iv) a user-friendly search interface based on an extensible catalogue of query templates in plain English. The main purpose of (iv) is to demonstrate that the different database models can be jointly queried based on our approach, but our system supports any 1.1 SPARQL-compliant general-purpose query builders, such as in (34–39).

Our article is structured as follows. In Section 2, we describe the individual databases, as well as a high-level introduction to our approach and the semantic models used in this work. In Section 3, we provide the implementation details of the three layers of our proposed architecture (data store, structured query interface, and application). In Section 4, we evaluate the performance of the system on a catalogue of 12 representative federated biological queries. Finally, we conclude with a discussion and outlook.

## 2. System and Methods

To understand more concretely the problem of integrating data from multiple sources, consider the following motivating example: “*What are the human genes which have a known association to glioblastoma (a type of brain cancer) and which furthermore have an orthologous gene expressed in the rat’s brain?”*. To answer this question, we would need to integrate information currently found in different databases:

1. Human proteins associated with glioblastoma can be obtained from **UniProt Knowledge Base**, a database providing a comprehensive, high-quality sequence and functional information on proteins (16). In the rest of the paper, we will use the name “UniProt” for readability.
2. The orthologs of these proteins in the rat can be obtained from **OMA** (Orthologous Matrix), a database of orthology inferences (30). Orthologs are genes in different species that evolved from a common ancestral gene by speciation. The orthologs are normally thought to retain the same function in the course of evolution. Other homology information, such as one-to-one orthology or paralogy, can be derived from the Hierarchical Orthologous Groups (HOGs) data structure (11, 40).
3. The genes expressed in the rat brain can be obtained from **Bgee**, a database of curated gene expression patterns in animals (31). Bgee version 14.0 includes gene expression data for 29 species such as human, mouse, or hedgehog. Currently, Bgee data are stored in a MySQL relational database (41).

In the following, we first provide a high-level description of our approach, then introduce the semantic models involved.

### 2.1. A federated, ontology-driven data integration approach

In order to achieve semantic interoperability between Bgee, OMA and UniProt, we have chosen a federated approach based on ontologies (Figure 1). The advantage of a federated approach is to avoid imposing a common global schema or meta-model on all data sources, and to facilitate the integration of further resources in the future. In doing so, we avoid, for example, the fastidious and time-consuming task of maintaining a centralised, integrated knowledge base. Instead, we provide a homogeneous data access layer to query the heterogeneous data sources. This homogeneous layer is part of a new generation of federated databases, such as poly-stores (42), that provide seamless access to distinct data models of storage engines (e.g. MySQL and RDF stores). Unlike that approach, we do not seek to optimise query performance by transferring data on-the-fly between disparate storage engines (42), but rather focus on solving syntactic and semantic heterogeneities among data stores.

**Fig. 1.**
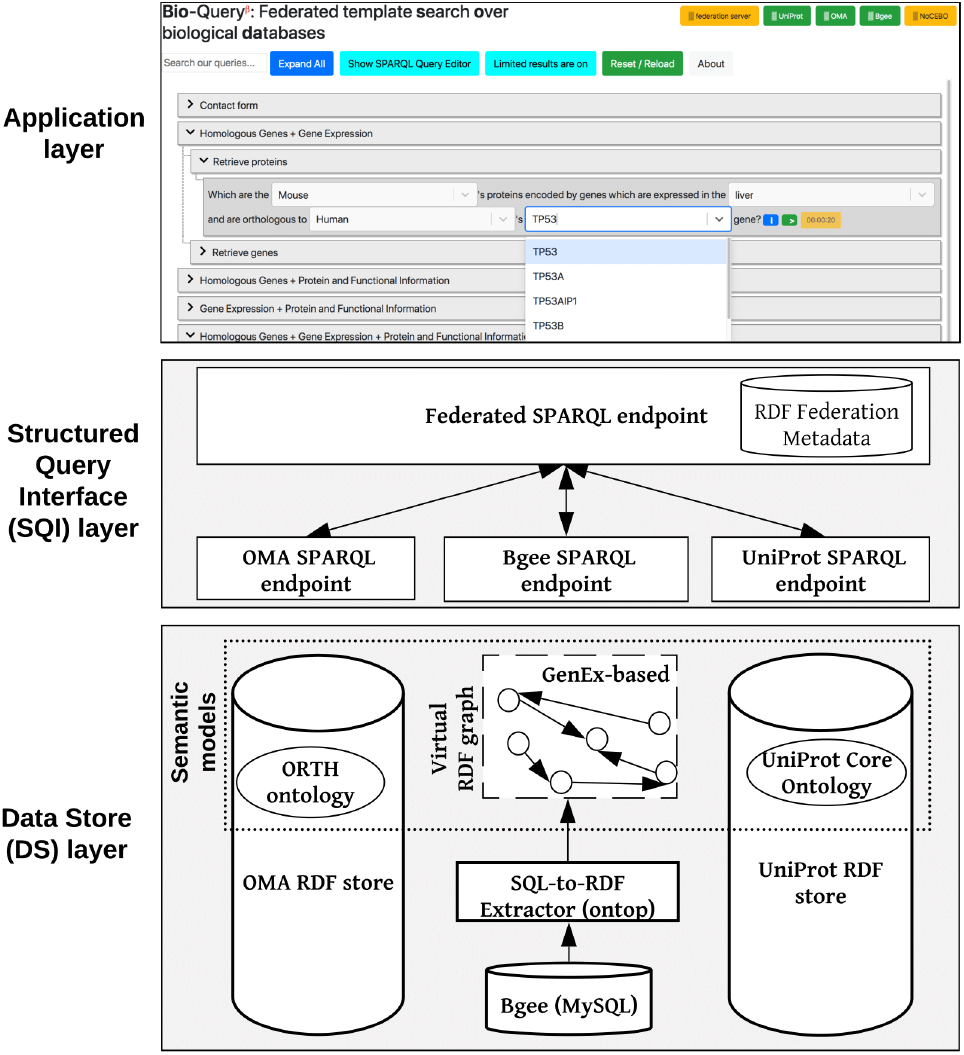
Overview of the ontology-driven federated data integration architecture applied to Bgee, OMA, and UniProt.The application layer depicts a Web search interface with editable templates to jointly query the data stores. Available online at http://biosoda.expasy.org.

To solve the syntactic heterogeneity, we rely on a structured query language—SPARQL—as the homogenous syntax to query all the data (32). We favoured SPARQL 1.1 over alternatives because it is World Wide Web Consortium (W3C) compliant and the data to be integrated are on the web; because it supports federated queries; and because one of our target data stores, UniProt, is already accessible through a SPARQL 1.1 endpoint, alongside a growing number of other biological databases (29). Indeed, although our initial prototype integrates data from Bgee, OMA and UniProt, we plan to extend the system to include more data sources in the future.

To reduce semantic heterogeneity among the databases, we rely on ontologies further described in Subsection 2.2, which are defined with the Web Ontology Language 2 (OWL 2), and thus based on the RDF model and syntax (https://www.w3.org/TR/owl2-overview/). RDF-based modelling decisions were taken to mitigate this heterogeneity when adopting ontological terms and instances to structure and represent the non-RDF data — OMA HDF5 and Bgee relational data. For example, by considering OMA, Bgee and UniProt databases, UniProt covers all of others with regard to the taxonomic lineage information for an organism. Therefore, we rely on UniProt classes and instance IRIs when representing and modelling taxonomy-related data in the OMA and Bgee RDF serialisations. To further exemplify, we can also mention the representation of genes among these three data sources. OMA genes completely overlaps Bgee genes but not all OMA genes has a corresponding one in UniProt, and *vice-versa*. Because of this, we decided to model Bgee genes the same way as in OMA, thereby easing interoperability between gene expression and orthology data. The next sections describe in more details the semantic models and the federated architecture proposed.

### 2.2. Semantic models

Ontology-based data integration requires as a preliminary step that each of the individual resources composing the federated system provide an explicit ontological description of their data. To minimise the need for semantic reconciliation—i.e. the process of identifying and resolving semantic conflicts (43), for example, by matching concepts from heterogeneous data sources (44)—we sought to rely as much as possible on existing ontologies when defining new semantic models.

Prior to our current work, among the three databases considered in this article, only UniProt provided an RDF representation of its data, as well as a SPARQL endpoint. The current UniProt RDF release comprises over 44 billion triples, and is based on the OWL 2 Full UniProt core ontology described in Redaschi et al. (45).

For the orthology data in **OMA**, we adopted the Orthology (ORTH) ontology (46), which was recently devised by the Quest for Orthologs Consortium (47) as a common data schema for integrating Orthology databases, such as OMA. We use ORTH to structure the OMA data, which is primarily stored in an HDF5 data store. Furthermore, during the conception of a second version of ORTH, design decisions such as the adoption of taxon-related terms from the UniProt ontology were made in order to enhance interoperability, enabling us to establish links among the data stores (Subsection 3.2.2). Therefore, the work presented in this article also contributed towards a new, improved version of the ORTH ontology, which is described in de Farias et al. (11).

In the case of **Bgee**, representing the original data in RDF proved to be a challenge, due to a general lack of a comprehensive ontology to serve as a data schema for describing knowledge in the field of gene expression. This may seem surprising considering the ubiquity of gene expression analyses in molecular biology and the existence of multiple well-established resources for gene expression—not only Bgee, but also Expression Atlas (EA) (48), Genevestigator (49), or the Tissue Expression database (50). We note here that different gene expression databases often use distinct criteria to assert *expressed in* or *absent in* relations.

To the best of our knowledge, two semantic models currently exist as initial attempts to structure gene expression related data: the Relation Ontology (51) and the Expression Atlas model (52). The Relation Ontology (RO) defines only a few terms within the domain of gene expression and is not specifically designed for this knowledge domain. Notably, it contains *expressed in* and *expresses* relations. The Expression Atlas defines a semantic model related to gene expression that mainly focuses on modelling the Expression Atlas (EA) data itself and not the domain of gene expression generally. In this EA model, additional data interpretations (i.e. semantics) are not explicitly represented, such as a given gene *is expressed* or *lowly expressed* in some sample relative to others. Although it would be possible to obtain this information through a more complex query on the Expression Atlas SPARQL endpoint, we lack an explicit representation, which would allow us to compare gene expression data from these different databases.

To provide a first step toward a general-purpose gene expression ontology, we drafted a new semantic model called GenEx. GenEx is aligned with the Relation Ontology and Expression Atlas models to facilitate interoperability with existing RDF stores. We also included semantic rules and terms to address (i) the representation of additional information related to gene expression, such as developmental stages, as well as *absent in* and *highly expressed* relations; and (ii) the trade-off between virtualisation and materialisation for the sake of query execution time and data storage. Furthermore, we reuse parts of the data schemas of the ORTH and UniProt core ontologies to provide (iii) the capacity to interoperate with other biological databases from different knowledge domains which are still relevant to the gene expression domain. For example, integrating orthology and gene expression data is relevant since we might want to predict gene expression conservation for orthologous genes. The draft GenEx is available online and documented in https://biosoda.github.io/genex/.

We stress that GenEx is currently in draft state. To become a standard, it needs to be endorsed and supported by multiple key stakeholders. We plan to initiate discussions with representatives of Bgee, Expression Atlas, Genevestigator, and Tissue database teams, and intend to solicit involvement from others, for example, the Model Organisms Databases (http://www.alliancegenome.org).

## 3. Implementation

Our federated data integration architecture comprises three layers: the data store (DS) layer, the structured query interface (SQI) layer, and the application layer (Figure 1). The DS layer contains all data stores to be integrated, including ontologies and methods to solve semantic and data model heterogeneities, such as relational-to-RDF mappings (Section 3.1). The SQI layer provides a homogeneous query language syntax and exploits common instances and literals (i.e. virtual links) to retrieve data from the DS layer (Section 3.2). The application layer includes any software tools that access the data stores through the SQI layer, for example a web search interface (Section 3.3). Figure 1 illustrates this architecture applied to our use case: the Bgee, OMA and UniProt databases.

The three layers are described in the next subsections, and source code is available at https://github.com/biosoda/bioquery.

### 3.1. Data store layer

The **UniProt** data were already available in an RDF model and accessible through a SPARQL endpoint at the start of our project. Therefore, we could use UniProt data as is.

The core of our work on the data store layer consisted in exposing data from **Bgee** and **OMA** as RDF, with the goal of solving data model heterogeneity. We focused our efforts on including the domain-specific, most “value-added” aspects of Bgee and OMA to the data store layer—leaving out information already available in UniProt. As a result, the Bgee and OMA data accessible through our system are subsets of their original contents. We provide an overview of the types of information available in the original sources versus in their RDF representation in the Supplementary Material (Table S3). This reduced the development work and data duplication among the databases, without loss of information considering that our federated approach enables directly retrieving this data from its original source (i.e. UniProt).

The **Bgee** data are stored in a relational database, meaning that integration between RDF stores and relational databases would still require substantial effort. There are two main methods to overcome this issue. First, the existing data could be represented entirely as RDF, which consequently would replace the relational model. A second approach would be to express the existing relational data as a virtual RDF graph, defined over ontological concepts and relations. We have chosen the latter approach, also referred to as *ontology based data access* (OBDA) (53). Our choice is justified by the fact that changing the Bgee data store into an RDF model would either lead to data duplication or would require significant changes in the current Bgee analysis pipeline (see 31). This is because Bgee is now adapted to the relational model for storing raw and preprocessed data from multiple data sources such as Ensembl, GEO, ArrayExpress and others (https://bgee.org/?page=source).

To implement OBDA over the Bgee relational database, we used the Ontop platform (53) version 3.0-beta-2. We defined several relational-to-RDF model mappings, which dynamically instantiate the gene expression semantic model described in Subsection 3.1. Figure 2 shows a simplified example of OBDA mappings that serve to express data from the relational model in the RDF model. Namespace prefixes such as *up:* shown in Figure 2 and used in the rest of this article are defined in Supplementary Table S1. While some of the mappings can be simple 1-to-1 correspondences—for example, a gene name (shown in red color on the right) can directly be used as a label of a *orth:Gene* class instance. Other mappings require transforming the original attributes in the relational data for interoperability – for example, replacing “:” with “_” in the case of anatomical entity identifiers from Bgee to be compliant with the existing UBERON ontology IRI (Internationalized resource identifier) terms (54), as shown in green color with the example of *UBERON:0000955* in Figure 2. Another type of transformation can be even combining multiple columns to instantiate a concept, as in the case of expressing *species* data from Bgee in terms of instances of *up:Taxon*. In this case, the OBDA mapping serves to concatenate the *genus* and *species* columns from Bgee in order to form the scientific name in compliance with the UniProt taxonomy. The scientific names of species in UniProt are denominated through the *up:scientificName* property, composed of both genus and species. This is illustrated in the left-most set of mappings (in blue color) in Figure 2. For further details of this OBDA mapping, see the Supplementary Material (Section S4). The full set of OBDA mappings used to expose Bgee relational data as virtual RDF triples are provided in https://github.com/biosoda/bioquery.

**Fig. 2.**
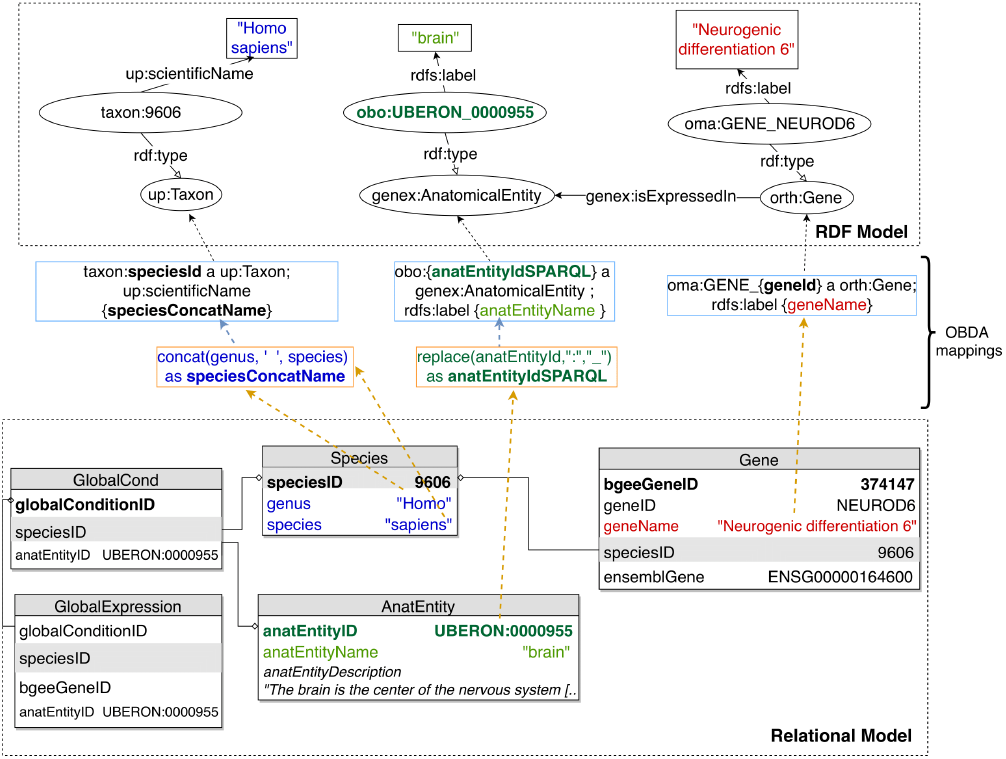
An illustration of relational-to-RDF mappings on a sample of the Bgee database. These mappings address both *schema-level* heterogeneity (an example is shown in blue), as well as *data-level* heterogeneity (shown in green). A mapping can also be a simple 1-to-1 correspondence between a relational attribute (e.g. *geneName*, shown in red) and its equivalent RDF property (in this case, an *rdfs:label* of an *orth:Gene* instance). Namespace prefixes are defined in Supplementary Table S1.

The code fragment in Listing 1 illustrates a mapping expressed with the Ontop relational-to-RDF mapping syntax, where the *source* is a SQL SELECT statement and the *target* consists of the corresponding RDF-based properties and classes. While direct and simple mappings (around 80% of the total) could in principle be automatically generated, complex ones such as the *isExpressedIn* relationship shown in Listing 1 can only be manually defined. Further explanations about this are available in Supplementary Material (Section S4).

**Listing 1.**
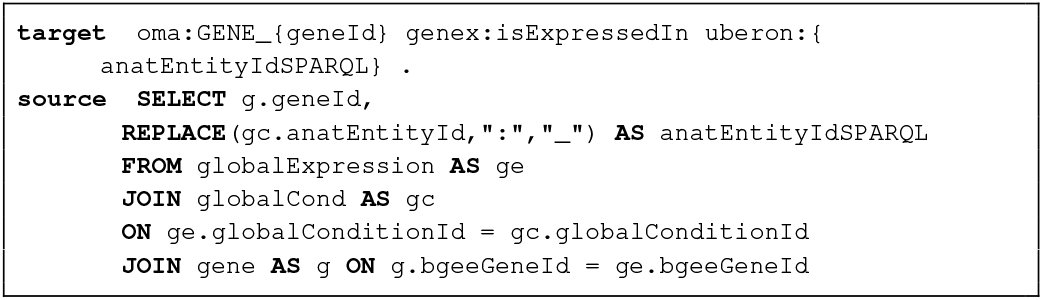
Ontop mapping to infer the “is expressed in” GenEx relation (i.e. target schema) based on the Bgee relational database (i.e. data source). Prefixes are defined in Supplementary Table S1.

Once relational-to-RDF mappings have been defined with Ontop between the Bgee MySQL database and GenEx, the original data can be queried with SPARQL, through the Bgee RDF virtual model. At query time, Ontop will translate SPARQL queries into SQL on-the-fly, using the mappings, and execute these over the Bgee relational database. On-top has the advantage of supporting federated queries as part of SPARQL 1.1 and of being open source. In order to enable researchers to directly use the RDF representation of Bgee, we made available a public SPARQL 1.1 endpoint at http://biosoda.expasy.org/rdf4j-server/repositories/bgeelight as a query service (without any webpage associated to it). Nonetheless, the OBDA solution with Ontop has some limitations – we discuss some of these in Section S4 in the Supplementary Material.

The **OMA** data are internally stored in an Hierarchical Data Format 5 (HDF5) file. This is not a database management system (DBMS) such as MySQL, but rather a data model and file format along with an API, libraries, and tools. Similarly to Bgee, we have to homogenise the OMA database model and syntax in order to enable integration with other biological RDF data stores (either virtual or materialised). For OMA, we chose to materialise the key parts of OMA data as an RDF graph, by implementing a hybrid approach, that combines materialisation and a possible RDF graph virtualisation for the sake of semantic enrichment and knowledge extraction, as described in detail in https://qfo.github.io/OrthologyOntology. The OMA RDF data and ORTH ontology are stored in a Virtuoso 7.2 triple store and a SPARQL endpoint is available at https://sparql.omabrowser.org/sparql. Further explanations regarding the OMA RDF data materialisation are available in Supplementary Material Section S5.

### 3.2. Structured query interface layer

Once the data stores are accessible through SPARQL endpoints, as depicted in Subsection 3.1, we can exploit means to link them at the data level. To do so, we identify common class instances and literals (e.g. strings) in order to establish “virtual links”. We define a virtual link as an intersection data point between two data stores. The links are required in order to enable performing federated queries, given that they act as join points between the federated sources. Figure 3 illustrates virtual links among UniProt, Bgee and OMA. For example, OMA and Bgee describe complementary information about common genes (instances of the *orth:Gene* class), as well as taxa (instances of the *up:Taxon* class), both of which can serve as virtual links to connect the two sources. A federated SPARQL query written based on the virtual links is described in Supplementary Material Section S3. To formally and explicitly describe virtual links, we adapted and extended the VoID RDF schema vocabulary (33) to include the concept of virtual links. We call this vocabulary Extended VoID (VoIDext). VoIDext is fully specified and exemplified in https://biosoda.github.io/voidext/. The entire metadata of virtual links among UniProt, Bgee, and OMA RDF stores for the work depicted in this article are available at http://purl.org/query/bioquery. In VoIDext specification, we also depict the SPARQL queries to retrieve the virtual links among OMA, Bgee and UniProt that support the writing of joint federated SPARQL queries. These queries can be executed on the federated SPARQL endpoint illustrated in Fig. 1 http://biosoda.expasy.org:8890/sparql. To leverage virtual links between Bgee and the other databases, we took advantage of the flexibility provided by Ontop when defining the Bgee OBDA mappings. We aimed at mapping Bgee data into corresponding instance IRIs and literals that already exist in OMA and UniProt RDF graphs. For example, species-related instance IRIs in the Bgee virtual graph are indeed exact matches of *up:Taxon* instance IRIs that are stored in the UniProt database.

**Fig. 3.**
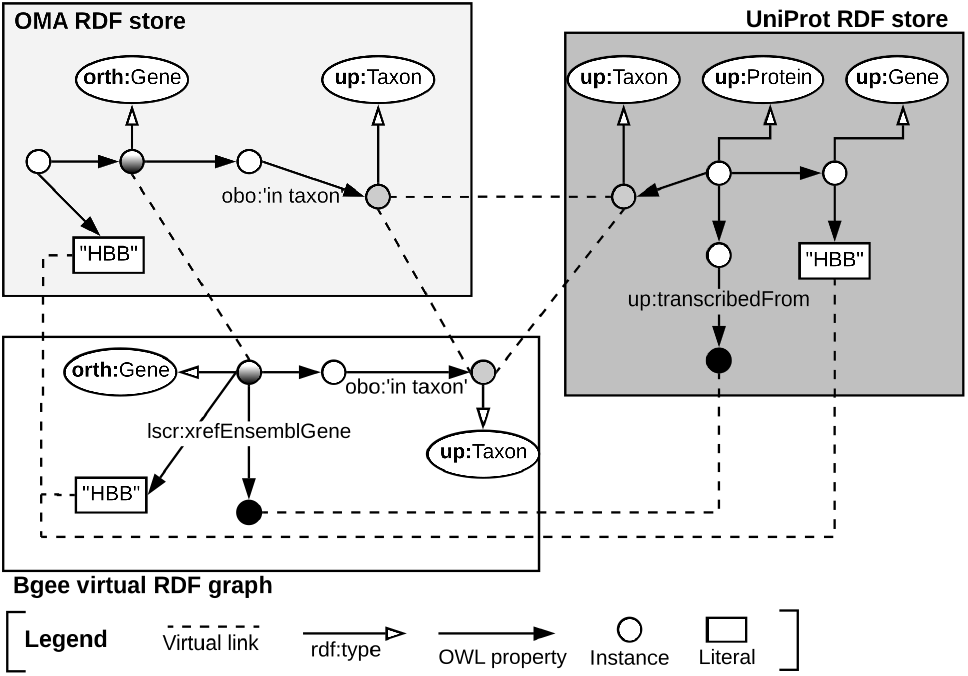
Example of virtual links among UniProt, OMA and Bgee data stores.

Likewise, the code fragment in Listing 1 asserts the (re)use of **OMA** gene instances as part of the **Bgee** virtual RDF graph rather than creating new Bgee ones. In this way, we avoid additional *owl:sameAs* assertions to state that the two instances are actually the same. Thus, *orth:Gene* instances are intersection nodes (i.e. virtual links) between the Bgee and OMA graphs. Figure 3 (left-hand side) illustrates a shared *orth:Gene* instance between OMA and Bgee graphs. Further information about the virtual links depicted in Figure 3 is available in Section S6 in the Supplementary Material.

Overall, we provide a federated SPARQL query endpoint along with an RDF store that exclusively contains metadata about the virtual links, and the SPARQL endpoints of the Uniprot, OMA and Bgee data stores. These metadata based on the VoIDext schema precisely define and document how the distributed datasets can be interlinked. Therefore, they may significantly facilitate the manual or automatic writing of a SPARQL 1.1 federated query, given that users are no longer required to discover the interlinks between the queried datasets on their own. In (55), we detail the drawbacks of the current VoID link sets to represent virtual links, and the description of the novel VoIDext specification.

### 3.3. Application layer

The main goal of the application layer in our work is to enable users, even with no prior technical training, easy access to the integrated information from the three biological databases. We developed a user-friendly interface (illustrated in the top part of Figure 1), which is accessible at http://biosoda.expasy.org/. The interface presents a catalogue of representative query templates drafted together with domain experts. The queries are provided in natural language, with editable fields, and grouped in a tree structure according to the target knowledge domain(s) and information retrieved for each query. A search bar is also provided, which enables filtering the templates by keywords of interest (e.g. “disease”).

For users with more advanced technical expertise, we also provide the option to show and modify the equivalent SPARQL queries. In doing so, our approach has the potential to increase the productivity of domain scientists in exploring the three heterogeneous data sets jointly. Additionally, the catalogue of questions is destined to grow according to user needs and feedback.

Moreover, because of the federated architecture of our system, its performance depends on that of the underlying data sources, e.g. UniProt. The availability of the underlying data stores is indicated by green labels in the top right corner of the webpage. For unavailable sources, the corresponding label is shown in yellow, as illustrated in the application layer in Figure 1.

By default, our system limits the total number of results returned, which allows for a faster response – the estimated response time is shown as a tag next to each query. However, the user can turn the limit option off, in order to obtain the full set of results. In this case, the response time may be significantly higher, which can largely be attributed to the SPARQL query execution time on the underlying sources. This has already been noted in similar previous systems (56). In terms of scalability, the UniProt SPARQL endpoint is a good example, having already an active user base of more than one thousand users per month. As we directly rely on the infrastructure of the underlying sources, we can therefore expect our system to exhibit reasonable performance for approximately the same number of users.

## 4. Results

In this section, we first revisit our motivating example for integrated data access to the three databases (UniProt, OMA and Bgee) and then present experimental results based on a catalogue of 12 federated queries. All results are reproducible through our public interface described in Subsection 3.3.

Recall our motivating example from the start of Section 2: “*What are the human genes which have a known association to glioblastoma (a type of brain cancer) and which furthermore have an orthologous gene expressed in the rat’s brain*”. Answering the question requires solving the following three subqueries:

1. Retrieve human proteins with a disease description related to glioblastoma from UniProt.
2. Retrieve orthologs of these proteins in the rat from OMA.
3. Only keep those orthologs for which there exists evidence of expression in the rat’s brain from Bgee.

The above steps translate to the federated SPARQL query in Supplementary Listing S2. The query produces a set of 15 human-rat orthologous pairs and can be executed in any SPARQL 1.1 endpoint. The full query, as well as a detailed list of results is available in the Supplementary Material in Section S3.

We further evaluated the performance, in terms of runtime, of 12 federated queries that illustrate real use cases requiring information across the three databases (see Table 1). The results are reproducible through our public template-based search interface. A detailed analysis of the queries, including their natural language description, the equivalent federated SPARQL queries, as well as an explanation of the complexity for each query, can be found at https://github.com/biosoda/bioquery.

**Table 1.** Descriptions of the 12 federated queries across OMA, Bgee, UniProt used for evaluating our system. The queries can be further refined and executed through our template search interface available at http://biosoda.expasy.org/.

| Query | Description |
| --- | --- |
| Q1 | Proteins in OMA encoded by the INS gene and their evidence type from UniProt. |
| Q2 | Rabbit proteins encoded by genes orthologous to the HBB-Y gene in Mouse and their associated information from UniProt. |
| Q3 | Rattus Norvegicus proteins paralogous to Tp53 and their UniProt function annotations. |
| Q4 | Mouse genes expressed in the liver which are orthologous to the human INS gene. |
| Q5 | Genes orthologous to a gene expressed in the fruit fly brain. |
| Q6 | Genes in Primates orthologous to a gene expressed in the fruit fly brain. |
| Q7 | Anatomic entities where the ins zebrafish gene is expressed and the ins gene GO annotations. |
| Q8 | Genes expressed in the human pancreas and their annotation in UniProt. |
| Q9 | Genes expressed in the human brain during the infant stage and their UniProt disease annotation. |
| Q10 | The orthologs of a gene that is expressed in the fruit fly brain and the UniProt annotations of these orthologs. |
| Q11 | The orthologs in **Primates** of a gene that is expressed in the fruit fly brain and the UniProt annotations of the primate orthologs. |
| Q12 | Proteins in humans, with a disease annotation, that are orthologous to a gene expressed in the rat brain. |

Table 2 shows that most of the queries can be executed in a few seconds – up to 6 seconds for 9 out of 12 queries, with less than half a second for 3 out of these. This holds even for queries with higher complexity (number of triple patterns). A triple pattern is similar to a regular RDF triple, except that any part of the triple can be replaced by a variable (32). Although preliminary, the results in Table 2 are encouraging for the use of SPARQL queries in data exploration tasks or in an interactive environment.

**Table 2.** Tests performed to evaluate our approach in terms of query execution time and the number of results. We evaluated 12 federated queries of varying complexity (measured in terms of number of triple patterns). Their description is provided in Table 1. All queries were executed twenty times, providing an average runtime and its standard deviation, given in seconds. The longest running query, Q10, is highlighted in bold.

| Query | Sources | #Results | Mean run-time (s) | Stddev run-time (s) |
| --- | --- | --- | --- | --- |
| Q1 | OMA, UniProt | 27 | 5.13 | 0.13 |
| Q2 | OMA, UniProt | 3 | 0.36 | 0.02 |
| Q3 | OMA, UniProt | 1 | 0.47 | 0.05 |
| Q4 | OMA, Bgee | 2 | 0.37 | 0.05 |
| Q5 | OMA, Bgee | 5322 | 4.72 | 0.2 |
| Q6 | OMA, Bgee | 38 | 2.38 | 0.09 |
| Q7 | Bgee, UniProt | 16 | 68.18 | 105.85 |
| Q8 | Bgee, UniProt | 58 | 33.17 | 25.23 |
| Q9 | Bgee, UniProt | 6 | 2.37 | 0.04 |
| **Q10** | **Bgee, OMA, UniProt** | **2269** | **349.18** | **4.19** |
| Q11 | Bgee, OMA, UniProt | 81 | 6.4 | 0.14 |
| Q12 | Bgee, OMA, UniProt | 3 | 5.24 | 0.11 |

The outlier Q10 calls for discussion. By comparing the natural language description of Q10 against Q11 (see corresponding entries in Table 1), we can intuitively deduce that the complexity stems from the high degree of generality of the sub-query that targets orthology information (OMA). In the case of Q10, retrieving an answer will require scanning the entire available orthology data and retrieving a large intermediate result set (orthologs found in any species, a total of 2269 results). By contrast, Q11 restricts the search space to the “primates” taxon only, which in practice results in a much lower query execution time (and a total of only 81 results). An important lesson derived from this example is that queries should always be as specific as possible, in order to limit both the search space and the size of intermediate results to the minimum necessary to obtain a relevant answer. Although this query illustrates a worst-case scenario, the results are still returned in less than 6 minutes—a latency which is tolerable for investigations in a biological research context.

## 5. Discussion and outlook

Data integration across heterogeneous biological databases promises to be one of the catalysts for gaining new biological insights in the postgenomic era. Here, we introduced an ontology-driven approach to bioinformatic resource integration. This approach enables complex federated queries across multiple domains of biological knowledge, such as gene expression and orthology, without requiring data duplication. The integration of the three sources promises to open the path for novel comparative studies across species, for example through the analysis of orthologs (OMA) of human disease-causing genes (UniProt) and their expression patterns in model organisms (Bgee). Thanks to modelling decisions made at the semantic (ontology) and data (assertions) levels, we established various virtual links among Bgee, OMA and UniProt data stores. Moreover, making these virtual links available in VoidExt facilitates the task of writing federated SPARQL queries, since users have an explicit representation of the connections (join points) between the three data sources. We furthermore lay the groundwork for bringing the benefits of integrated data to domain specialists through a template-based search engine available online, which does not require users to know SPARQL in order to pose questions on the integrated data.

The catalogue of federated queries across the three data sources can serve as a starting point towards answering new biological questions that span across the domains of evolutionary relationships and gene expression. The results presented in this study can be easily reproduced through our template search interface. We furthermore make available all source code, including the template search interface code, relational-to-RDF mappings and the catalogue of queries, with the goal of facilitating reuse of these components for further research. All resources are available in our GitHub repository.

Our experiments show that most queries in our catalogue can be answered within seconds. And although the more complex queries take several minutes to complete, we expect this turnaround time to be tolerable for most interested users— particularly considering the alternative of manually querying the resources and combining the results. In the future, we plan to add a federated query optimiser to our system to further improve the response time. We also note here that the application interface directly queries the underlying databases without performing additional tasks, such as considering all gene name synonyms to get broader results. We plan to support such features as part of future work. To support virtual link evolution, we aim to develop a tool to automatically detect broken virtual links because of either data schema changes or radical modifications of instances’ IRIs and property assertions. Meanwhile, we encourage contributions to the current query catalogue, which will serve in the study of a natural language search interface for the integrated biological data as part of future work.

## ACKNOWLEDGEMENTS

This work was supported by the Swiss National Research Programme 75 “Big Data” [167149], the Swiss National Science Foundation [150654], and the Swiss State Secretariat for Education, Research and Innovation (SIB Infrastructure grants).

