## Supplementary Material for "Enabling Semantic Queries Across Federated Bioinformatics Databases"

### S1. Example that compares the number of UniProt protein entries between the Linked Life Data and UniProt RDF stores

The following example illustrates one of the main drawbacks of centralized data integration approaches, namely that in the lack of a good strategy for keeping data in-sync, results can quickly become stale. As an illustration, we can compare the total number of protein entries in the Linked Life Data (LLD) SPARQL endpoint ((1) mentioned in the Introduction section in the paper), with the count according to the latest version of UniProt.
Executing the SPARQL query in the version of UniProt at the time of writing (release 2019_03) will result in a total of around 238 million proteins, whereas in Linked Life Data there are only around 20 million, indicating that the entries in LLD are severely outdated, missing out more than 90% of the total entries in the latest UniProt version. We provide below the required links in order to reproduce these observations:
Query to retrieve the total number of proteins:

```
PREFIX up:<http://purl.uniprot.org/core/>
SELECT (count(?protein) as ?count_uniprot_entries )
WHERE
{ ?protein a up:Protein .}
```

**Executing the query at Linked Life Data SPARQL endpoint**:
http://www.linkedlifedata.com/sparql

To get the list of results in your browser directly, use the following link:
http://www.linkedlifedata.com/sparql?query=PREFIX+up%3A%3Chttp%3A%2F%2Fpurl.uniprot.org%2Fcore%2F%3E+%0D%0ASELECT+%28count%28%3Fprotein%29+as+%3Fcount_uniprot_entries+%29%0D%0AWHERE%0D%0A%7B%0D%0A%09%3Fprotein+a+up%3AProtein+.%09%0D%0A%7D&_implicit=false&implicit=true&_form=%2Fsparql

**Executing the query at UniProt SPARQL endpoint**:
https://sparql.uniprot.org/sparql/

To get the list of results in your browser directly, use the following link:
https://sparql.uniprot.org/sparql/?format=html&query=PREFIX+up%3A%3Chttp%3A%2F%2Fpurl.uniprot.org%2Fcore%2F%3E+%0D%0ASELECT+%28count%28%3Fprotein%29+as+%3Fcount_uniprot_entries+%29%0D%0AWHERE%0D%0A%7B%0D%0A%09%3Fprotein+a+up%3AProtein+.%09%0D%0A%7D

### S2. An example of Bgee-Ontop relational-to-RDF mapping for the species table

The code fragment in Listing S1 defines how rows (i.e. a species) and columns (i.e. species attributes) of the *species* table in the relational database can be mapped as RDF triples, by instantiating a corresponding GenEx class, namely the *up:Taxon* class (imported from the UniProt core ontology), as well as its individuals. The mapping also addresses *schema-level heterogeneity* by concatenating two source column values (i.e. *genus* and *species*) into the target *up:scientificName* RDF property.

```
target   taxon:{speciesId} a up:Taxon ;
              up:scientificName {speciesSciName} ;
              up:rank  up:Species ;
              up:commonName {speciesCommonName}.
              orth:Organism/{speciesId} a orth:Organism ;
              obo:RO_0002162 taxon:{speciesId} .
source   SELECT speciesId, speciesCommonName,
         CONCAT(genus, '␣', species) AS speciesSciName
         FROM species
```

**Listing S1.** Ontop direct mapping to infer Species related data into the GenEx semantic model (i.e. target schema) based on the Bgee relational database (i.e. data source). Prefixes are defined in Table S1.

**Table S1.** In this article, we assume the namespace prefix bindings in this table.

| Prefix | Namespace Internationalized Resource Identifier IRI |
|---|---|
| rdfs: | http://www.w3.org/2000/01/rdf-schema# |
| rdf: | http://www.w3.org/1999/02/22-rdf-syntax-ns# |
| orth: | http://purl.org/net/orth# |
| up: | http://purl.uniprot.org/core/ |
| taxon: | http://purl.uniprot.org/taxonomy/ |
| genex: | http://purl.org/genex# |
| obo:, uberon: | http://purl.obolibrary.org/obo/ |
| oma: | http://omabrowser.org/ontology/oma# |
| skos: | http://www.w3.org/2004/02/skos/core# |
| sio: | http://semanticscience.org/resource/ |
| lscr: | http://purl.org/lscr# |
| void: | http://rdfs.org/ns/void# |
| voidext: | http://purl.org/query/voidext# |

### S3. Example SPARQL federated query across Bgee, OMA, UniProt

*What are the human genes which have a known association to glioblastoma (a type of brain cancer) and which furthermore have an orthologous gene expressed in the rat brain?*

```
PREFIX rdfs: <http://www.w3.org/2000/01/rdf-schema#>
PREFIX obo: <http://purl.obolibrary.org/obo/>
PREFIX orth: <http://purl.org/net/orth#>
PREFIX sio: <http://semanticscience.org/resource/>
PREFIX taxon: <http://purl.uniprot.org/taxonomy/>
PREFIX up: <http://purl.uniprot.org/core/>
PREFIX lscr: <http://purl.org/lscr#>
PREFIX genex:<http://purl.org/genex#>

SELECT DISTINCT ?protein ?orthologous_protein_rat ?id WHERE {
 SELECT * {
  SERVICE <http://sparql.uniprot.org/sparql> {
    SELECT ?protein WHERE {
      ?protein a up:Protein ;
               up:organism taxon:9606 ;
               up:annotation ?annotation .
      ?annotation rdfs:comment ?annotation_text .
      ?annotation a up:Disease_Annotation .
    FILTER CONTAINS (?annotation_text, "glioblastoma") } }
  SERVICE <https://sparql.omabrowser.org/sparql> {
    SELECT ?orthologous_protein_rat ?protein ?id WHERE {
      ?protein_OMA a orth:Protein .
      ?orthologous_protein_rat a orth:Protein .
      ?cluster a orth:OrthologsCluster .
      ?cluster orth:hasHomologousMember ?node1 .
      ?cluster orth:hasHomologousMember ?node2 .
      ?node2 orth:hasHomologousMember* ?protein_OMA .
      ?node1 orth:hasHomologousMember* ?orthologous_protein_rat .
      ?orthologous_protein_rat orth:organism/obo:RO_0002162 taxon:10116. #rattus norvegicus
      ?orthologous_protein_rat sio:SIO_010079/lscr:xrefEnsemblGene ?id .
      ?protein_OMA lscr:xrefUniprot ?protein .
      FILTER(?node1 != ?node2) } }
  SERVICE <http://biosoda.expasy.org:8080/rdf4j-server/repositories/bgeelight> {
                ?gene genex:isExpressedIn ?a .
                ?a rdfs:label "brain" .
                ?gene orth:organism ?s .
                ?s obo:RO_0002162 taxon:10116.
                ?gene lscr:xrefEnsemblGene ?id . } } }
```

**Listing S2.** A federated SPARQL 1.1 query to retrieve proteins associated with glioblastoma and the orthologs expressed in the rat brain.

#### S3.1. Results

Table S2 displays the results of executing the SPARQL query above, where:

- The first column, “protein”, shows UniProt human proteins with a known association with glioblastoma for which there exists an orthologous protein expressed in the rat’s brain. Clicking on any of the links in this column will redirect to the corresponding UniProt entry online.
- The second column, “orthologous_protein_rat”, shows the orthologous rat protein (for which there exists known expression in the brain according to data from Bgee)
- The third column, “id”, shows the Ensembl ID of the gene encoded by the rat protein (from column 2). Note that the ensemble ID (e.g. ENSRNOG00000008839) can be used in the Bgee search interface at https://bgee.org/ for validating the results.

The complete list of federated queries is available in our template-based search interface at http://biosoda.expasy.org, while an explanation for the complexity of the queries is available in our github repository at https://github.com/biosoda/bioquery/.

**Table S2.** Results of federated SPARQL query joining Bgee, OMA and UniProt.

| protein | orthologous_protein_rat | id |
|---|---|---|
| http://purl.uniprot.org/uniprot/P37231 | https://omabrowser.org/oma/info/RATNO15188 | http://rdf.ebi.ac.uk/resource/ensembl/ENSRNOG00000008839 |
| http://purl.uniprot.org/uniprot/P08922 | https://omabrowser.org/oma/info/RATNO12308 | http://rdf.ebi.ac.uk/resource/ensembl/ENSRNOG00000000406 |
| http://purl.uniprot.org/uniprot/P68431 | https://omabrowser.org/oma/info/RATNO09038 | http://rdf.ebi.ac.uk/resource/ensembl/ENSRNOG00000053155 |
| http://purl.uniprot.org/uniprot/P68431 | https://omabrowser.org/oma/info/RATNO09042 | http://rdf.ebi.ac.uk/resource/ensembl/ENSRNOG00000056281 |
| http://purl.uniprot.org/uniprot/P68431 | http://omabrowser.org/oma/info/RATNO11352 | http://rdf.ebi.ac.uk/resource/ensembl/ENSRNOG00000060366 |
| http://purl.uniprot.org/uniprot/Q14956 | https://omabrowser.org/oma/info/RATNO14717 | http://rdf.ebi.ac.uk/resource/ensembl/ENSRNOG00000008816 |
| http://purl.uniprot.org/uniprot/P84243 | https://omabrowser.org/oma/info/RATNO18582 | http://rdf.ebi.ac.uk/resource/ensembl/ENSRNOG00000032401 |
| http://purl.uniprot.org/uniprot/P84243 | https://omabrowser.org/oma/info/RATNO06508 | http://rdf.ebi.ac.uk/resource/ensembl/ENSRNOG00000003220 |
| http://purl.uniprot.org/uniprot/O75140 | https://omabrowser.org/oma/info/RATNO07117 | http://rdf.ebi.ac.uk/resource/ensembl/ENSRNOG00000018144 |
| http://purl.uniprot.org/uniprot/Q12980 | https://omabrowser.org/oma/info/RATNO03263 | http://rdf.ebi.ac.uk/resource/ensembl/ENSRNOG00000020541 |
| http://purl.uniprot.org/uniprot/Q8WTW4 | https://omabrowser.org/oma/info/RATNO20234 | http://rdf.ebi.ac.uk/resource/ensembl/ENSRNOG00000021660 |
| http://purl.uniprot.org/uniprot/Q9BZH6 | https://omabrowser.org/oma/info/RATNO02052 | http://rdf.ebi.ac.uk/resource/ensembl/ENSRNOG00000020430 |
| http://purl.uniprot.org/uniprot/Q9HD26 | https://omabrowser.org/oma/info/RATNO12311 | http://rdf.ebi.ac.uk/resource/ensembl/ENSRNOG00000000408 |
| http://purl.uniprot.org/uniprot/Q9Y243 | https://omabrowser.org/oma/info/RATNO06482 | http://rdf.ebi.ac.uk/resource/ensembl/ENSRNOG00000021497 |
| http://purl.uniprot.org/uniprot/Q9UM73 | https://omabrowser.org/oma/info/RATNO17047 | http://rdf.ebi.ac.uk/resource/ensembl/ENSRNOG00000008683 |

### S4. Example relational-to-RDF OBDA mappings and discussion

Figure S1 – provided here as a copy of Figure 2 in the paper, for readability purposes – illustrates graphically an example of exposing relational data from Bgee (shown in the bottom half of the figure, the relational model) in a virtual RDF graph (a fraction of the ontology and 2 instances are shown in the upper half of the figure, the RDF model).

The orange rectangles in the OBDA mappings layer (in the center of the figure) represent the "source" fragments (a simplified SQL statement) of a relational-to-RDF mapping, while the blue rectangles illustrate the "target" (resulting RDF triples). For readability purposes, we only included 3 sample mappings in this figure. However, the full set of OBDA mappings employed

with the Bgee relational data is available in our github repository at https://github.com/biosoda/bioquery/tree/master/Bgee_OBDA_mappings.
We note here that more complex mappings OBDA mappings can require joining multiple tables in order to express an ontological concept or property which does not have a direct correspondent (i.e., is *not* explicitly present) in the original database. For this reason, mappings such as the one in Listing 1 from the paper (shown here in Listing S3 below) can be interpreted as a *semantic enrichment* of the Bgee original data schema, because they provide an explicit representation for knowledge that is not directly present in the base data. In addition, all RDF triples are indeed virtual, not materialised, hence the data are not duplicated.

```
target  oma:GENE_{geneId} genex:isExpressedIn uberon:{anatEntityIdSPARQL} .
source  SELECT g.geneId,
       REPLACE(gc.anatEntityId,":","_") AS anatEntityIdSPARQL
       FROM globalExpression AS ge
       JOIN globalCond AS gc
       ON ge.globalConditionId = gc.globalConditionId
       JOIN gene AS g ON g.bgeeGeneId = ge.bgeeGeneId
```

**Listing S3.** Ontop mapping to infer the "is expressed in" GenEx relation (i.e. target schema) based on the Bgee relational database (i.e. data source). Prefixes are defined in Supplementary Table S1.

In more detail, the code fragment in Listing S3 asserts the *genex:isExpressedIn* property by relating the projected columns *geneId* and *anatEntityIdSPARQL* from the join between the *globalExpression*, *globalCond* and *gene* tables in the Bgee database. This mapping also addresses *data-level heterogeneity* by applying the SQL REPLACE() function in order to transform the *anatEntityId* attribute (represented in Bgee with the separator ":") into the corresponding standard UBERON IRI (where the "_" separator is used). This transformation is also graphically illustrated in Figure S1 (in green).
Nonetheless, the OBDA solution with Ontop has some limitations. In particular, Ontop may struggle to apply complex and numerous mappings. In principle, the advantage of Ontop is that, by translating SPARQL queries into SQL, the system can take full advantage of the SQL query optimizer provided by the underlying relational database management system (RDBMS). However, when the relational-to-RDF mappings are very complex, the translation of a given SPARQL query can result in an extremely complex SQL equivalent that can possibly imply an overall poor performance of the system (Ontop + RDBMS) (2). Moreover, currently not all SPARQL queries can be translated into SQL. For example, aggregation queries (e.g. SUM, COUNT, MAX) are not yet supported.

### S5. Information available in Bgee, OMA and UniProt

In the Table S3 we provide an overview of the type of information available in Bgee, OMA and UniProt, both in the original representation (relational database for Bgee and HDF5 for OMA), as well as in RDF. More precisely, an "x" represents information available; an added "+" symbolizes information available in RDF; and "o" represents a link to other databases (e.g. OMA homologous groups).

**Table S3.** The information available on Bgee, OMA and UniProt data stores by also including resulted information (i.e. non-stored) after some data processing. Legend: "x" represents information available; "+" information available in RDF; and "o" represents a link to other databases (e.g. OMA homologous groups).

| Information | Bgee | OMA | UniProt |
|---|---|---|---|
| Gene Ontology annotations | x | x | x+ |
| Cross-references | x+ | x+ | x+ |
| Family and domain | x | x+ | |
| Local synteny | x | | |
| Pairwise homologous genes/proteins | x+ | | |
| Homologous groups of genes/proteins | o | x+ | o |
| Hierarchical Orthologous Group (HOG) | x+ | o | |
| Gene expression | x+ | o | |
| Absence of gene expression | x | | |
| Anatomical entity annotations (UBERON) | x+ | | |
| Developmental stage annotations | x+ | | |
| Species taxonomy (NCBI identifiers) | x+ | x+ | x+ (fully) |
| Gene and/or protein names | x+ | x+ | x+ |
| Subcellular location (including GO annotation) | x | x+ | |
| Sequences | x | x+ | |
| Post-translational modifications and/or processing events | x+ | | |
| Protein structures (quaternary, tertiary and secondary) | x+ | | |
| Similar proteins based on their membership in UniProt Reference Clusters (UniRef). | x+ | | |

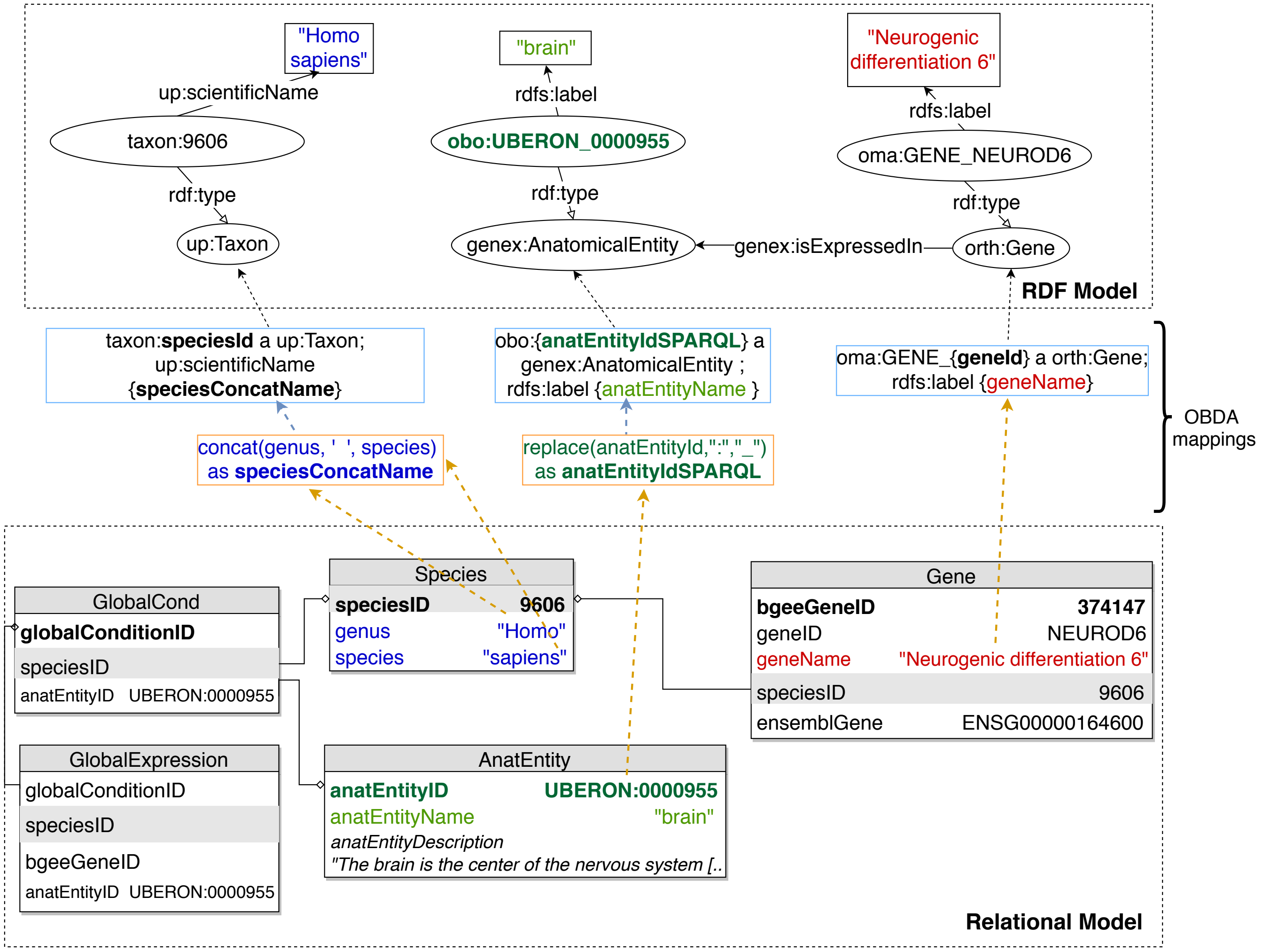


**Fig. S1.** An illustration of relational-to-RDF mappings on a sample of the Bgee database (a copy of Figure 2 in the paper, provided here for readability). These mappings address both *schema-level* heterogeneity (an example is shown in blue), as well as *data-level* heterogeneity (shown in green). A mapping can also be a simple 1-to-1 correspondence between a relational attribute (e.g. *geneName*, shown in red) and its equivalent RDF property (in this case, an *rdfs:label* of an *orth:Gene* instance). Namespace prefixes are defined in Supplementary Table S1.

#### S5.1. Discussion regarding choice of RDF materialization for OMA data

From the original OMA data, we chose to exclusively materialise in RDF the Hierarchical Orthologous Groups (HOGs) and the cross-references to other resources, which are the main "value-added" parts of OMA. The RDF serialisation relies on the ORTH ontology documented in `https://qfo.github.io/OrthologyOntology/`. Other information, such as pairwise orthologs (i.e. orth:hasOrtholog property assertions), may be inferred from the HOGs, therefore it does not require to be materialised (3).

The choice of a partial materialisation in the case of OMA is further justified by the following reasons:

(i) To our knowledge, there is currently no OBDA solution to query HDF5 data stores as virtual RDF graphs with SPARQL support. One possible direction would be to translate SPARQL into HDFql (i.e. a recent SQL-like query language for HDF5 data). In analogy to Ontop for Bgee, using such an approach would allow us to expose OMA data as an RDF virtual graph and convert the data to RDF on-the-fly. However, building an OBDA solution for HDF5 from scratch requires substantial development efforts, well beyond the scope of this project.

(ii) Parts of the data in the OMA HDF5 store are already available in UniProt (e.g. the protein sequences or the gene ontology annotations—for further examples see Supplementary Material). Hence, by solely materialising into RDF the OMA HDF5 data which are not already available through the UniProt SPARQL endpoint, we reduce the amount of data duplication, as well as the maintenance efforts required to keep data in sync with changes in UniProt.

### S6. Discussion regarding Virtual Links

At a technical level, note that virtual links among OMA, Bgee and UniProt can be classified into two different types: (i) IRI based (e.g. an instance), and (ii) data literal based (e.g. a label, such as "Homo sapiens" or "HBB"). For example, cross-reference IRIs serve as virtual links of type (i) among the data stores. Concretely, let us consider cross-references to Ensembl genes (4). On the one hand, the Ensembl gene IRI is assigned to the *up:transcribedFrom* OWL object property in the UniProt RDF store. This IRI is illustrated as a filled black circle in Figure 3. On the other hand, we reuse the same IRI as value of the *lscr:xrefEnsemblGene* OWL annotation property in the Bgee and OMA RDF graphs. To illustrate type (ii), we can mention the *rdfs:label* and *skos:prefLabel* properties that assert the same gene names (e.g. "HBB") to an instance of *orth:Gene* in the Bgee graph and *up:Gene* in the UniProt RDF graph, respectively. In Figure 3 in the paper, these literals are represented as rectangles.

1. Vassil Momtchev, Deyan Peychev, Todor Primov, and Georgi Georgiev. Expanding the pathway and interaction knowledge in linked life data. *Proc. of International Semantic Web Challenge*, 2009.
2. D Calvanese, B Cogrel, S Komla-Ebri, and others. Ontop: Answering SPARQL queries over relational databases. *Semant. Pragmat.*, 2017.
3. Tarcisio M de Farias, Hirokazu Chiba, and Jesualdo T Fernández-Breis. Leveraging logical rules for efficacious representation of large orthology datasets. April 2017.
4. Daniel R Zerbino, Premanand Achuthan, Wasiu Akanni, M Ridwan Amode, Daniel Barrell, Jyothish Bhai, Konstantinos Billis, Carla Cummins, Astrid Gall, Carlos García Girón, Laurent Gil, Leo Gordon, Leanne Haggerty, Erin Haskell, Thibaut Hourlier, Osagie G Izuogu, Sophie H Janacek, Thomas Juettemann, Jimmy Kiang To, Matthew R Laird, Ilias Lavidas, Zhicheng Liu, Jane E Loveland, Thomas Maurel, William McLaren, Benjamin Moore, Jonathan Mudge, Daniel N Murphy, Victoria Newman, Michael Nuhn, Denye Ogeh, Chuang Kee Ong, Anne Parker, Mateus Patricio, Harpreet Singh Riat, Helen Schuilenburg, Dan Sheppard, Helen Sparrow, Kieron Taylor, Anja Thormann, Alessandro Vullo, Brandon Walts, Amonida Zadissa, Adam Frankish, Sarah E Hunt, Myrto Kostadima, Nicholas Langridge, Fergal J Martin, Matthieu Muffato, Emily Perry, Magali Ruffier, Dan M Staines, Stephen J Trevanion, Bronwen L Aken, Fiona Cunningham, Andrew Yates, and Paul Flicek. Ensembl 2018. *Nucleic Acids Res.*, 46(D1):D754–D761, January 2018.
